# Establishment and characterization of a tumoroid biobank derived from dog patients’ mammary tumors for translational research

**DOI:** 10.1101/2022.09.04.506533

**Authors:** Antonella Raffo-Romero, Soulaimane Aboulouard, Emmanuel Bouchaert, Agata Rybicka, Dominique Tierny, Nawale Hajjaji, Isabelle Fournier, Michel Salzet, Marie Duhamel

**Affiliations:** Université Lille, Inserm, CHU Lille, U1192, Laboratoire Protéomique, Réponse Inflammatoire et Spectrométrie de Masse (PRISM), Lille, France; OCR (Oncovet Clinical Research), Parc Eurasanté Lille Métropole, 80 rue du Dr Yersin, 59120 Loos, France; Breast Cancer Unit, Oscar Lambret Center, Lille, France; Institut Universitaire de France, Paris, France

**Keywords:** breast cancer, dog patients, tumoroid, biobank, drug screening

## Abstract

Breast cancer is the most frequent cancer among women causing the greatest number of cancer-related deaths. Cancer heterogeneity is a main obstacle to therapies. Around 96% of the drugs fail from discovery to the clinical trial phase probably because of the current unreliable preclinical models. New models emerge such as companion dogs who develop spontaneous mammary tumors resembling human breast cancer in many clinical and molecular aspects. The present work aimed at developing a robust canine mammary tumor model in the form of tumoroids which recapitulate the tumor diversity and heterogeneity. We conducted a complete characterization of these canine mammary tumoroids through histologic, molecular and proteomic analysis, demonstrating their strong similarity to the primary tumor. We demonstrated that these tumoroids can be used as a drug screening model. In fact, we showed that Paclitaxel, a human chemotherapeutic, could killed canine tumoroids with the same efficacy as human tumoroids with 0.1 to 1 μM of drug needed to kill 50% of the cells. Due to easy tissue availability, canine tumoroids can be produced at larger scale and cryopreserved to constitute a biobank. We have demonstrated that cryopreserved tumoroids keep the same histologic and molecular features (ER, PR and HER2 expression) as fresh tumoroids. Two techniques of cryopreservation were compared demonstrating that tumoroids made from frozen tumor material allowed to maintain a higher molecular diversity. These findings revealed that canine mammary tumoroids can be easily generated at large scale and can represent a more reliable preclinical model to investigate tumorigenesis mechanisms and develop new treatments for both veterinary and human medicine.

## 1 Introduction

A major obstacle in preclinical drug development for cancer is the lack of appropriate cell culture model systems. Two-dimensional cancer cell lines are frequently used for the first screening of newly developed drugs and for the study of cancer development. However, cancer cell lines completely lack interaction with the tumor microenvironment, which is the main reason for drug resistance. Mouse models present also several drawbacks leading to difficulties in the translation to human diseases. Such models do not fully recapitulate the diversity and architecture of the primary disease, thus providing inaccurate analysis of tumor pathogenesis and sensitivity to therapy. Around 96% of the drugs fail from discovery to the clinical trial phase, probably because the preclinical models are not close enough to the tumor biology in patients(1).

Tumoroid cultures represent a robust three-dimensional (3D) *in vitro* system that faithfully recapitulate the genetic and phenotypic characteristics of the tumor from which they are derived. The 3D tumoroid system has been utilized to study different types of cancers(2–5). Tumoroids can serve to better understand the biology but also to test drug efficacy *in vitro* before clinical trials in human patients. Most of the tumoroid studies have been conducted on mouse and human tissue samples. Mouse tumor tissues do not fully recapitulate the human disease and therefore are not the best models for human translation. On the other hand, the use of human samples is the optimal solution but the difficulty to access to the fresh tissues and ethical issues can slow down the large scale screening of new drugs. That is why it is of prime importance that new models emerge than can fully and faithfully recapitulate the human disease. Moreover, as tumoroids are found to have more and more relevance and applications, large scale production of tumoroids become inevitable but presents many challenges due to the difficulty of accessing large quantities of human fresh tissue.

In that regard, companion dogs with spontaneous tumors present a unique, ethical, non-experimental model for comparative research and drugs development(6,7). Canine mammary tumor (CMT) is the third most common type of cancer in dogs, and first in bitches with an incidence of around 230 cases per 100,000 dogs per year(8–10). It possesses several advantages over highly inbred and genetically modified laboratory animals, such as clinical profile (age at onset, predominance of carcinomas, and type of metastases), genetics (role of BRCA1/2, overlapping gene signature) and molecular similarities with its human counterpart(11,12). CMT, the same as human breast cancer (BC), can be characterized by expression of estrogen, progesterone and HER2 receptors. The involvement of companion dogs with spontaneous CMT in translational oncology is already seen in numerous publications and several ongoing clinical trials(13). Canine tumoroids developed from dog patients with spontaneous CMT could therefore provide a more representative and ethical translational model to test drug efficacy and toxicity in pre-human studies, as well as canine tumoroids could be an innovative screening tool in drug discovery, while reducing the number of experimental animals needed for *in vivo* studies. Few canine tumoroids studies have been made so far(14–17), one of them has developed tumoroids from canine normal and tumor breast stem cells(18). None of them have developed tumoroids from CMT heterogeneous tissue recapitulating the tumor microenvironment.

The enormous potential of tumoroids as preclinical models has given rise to the development of tumoroid biobanks. Tumoroid biobanks have been obtained from various tumor tissues(19). However, there is a lack of knowledge about cryopreservation procedures and whether the cryopreserved tumoroids maintain similar molecular and functional features as their fresh counterparts.

In this study, we have developed for the first time tumoroids from heterogeneous canine mammary tumor tissues. We have demonstrated morphologic, histologic and molecular stability between fresh CMT tumoroids and cryopreserved tumoroids. However, tumoroids made from frozen CMT material show more similarities with fresh tumoroids compared to cryopreserved tumoroids. Treatment with a chemotherapeutic drug also confirmed these results. Taken together, our study aimed at creating and characterizing a new biobank of canine mammary tumoroids with similar features as human tissues which can be used for large scale drug screening in preclinical studies.

## 2 Materials and Methods

### 2.1. Human and Dog patients’ tissue collection

This study was carried out with canine mammary tumors (n = 6). The tumors were collected at different veterinary clinics from dogs undergoing scheduled surgery. The samples were delivered with the written consent of the owners. The dogs included in the study were treated surgically by their veterinarian, and none of them received any additional treatment before the mastectomy. A veterinary pathologist reviewed the tissue blocks to confirm the diagnosis and define the lesions for dissection. For this study, we received a piece of fresh tumor of approximately 1 cm3.

Human breast tumor tissue was obtained from a patient undergoing surgery for early breast cancer. Fresh tumor tissue was provided by the pathologist for organoid culture. The sample was anonymized prior to its transfer to the lab. The study was approved by the local research committee of Oscar Lambret Cancer center and a French Ethical Committee (study IdRCB 2021-A00670-41). The written informed consent for the study was obtained from the patient before any procedure.

### 2.2. Tissue processing

Each tumor sample was divided into three pieces: one piece was snap frozen in liquid nitrogen before being stored at −80°C for proteomics large scale study, the second piece was fixed in formaldehyde 4% for 24h followed by dehydration in 20% sucrose for 24h, embedding in gelatin and storage at −80°C for histopathological analysis and hematoxylin and eosin staining. The last tumor fragment was used for tumoroids culture. For this, the tumor fragment was minced into 1 mm3 pieces before its enzymatic digestion as described below.

### 2.3. Tumoroid culture

The minced tumor tissue was digested in 2 mL Hank’s Balanced Salt Solution (HBSS, Gibco) with antibiotics and anti-fungal (1X Penicillin/Streptomycin, 1X Amphoteromicin) containing 1 mg/mL collagenase type IV (Sigma) and 5 U/ mL hyaluronidase (Sigma) at 37 °C for 2 h. During this time, the medium containing the tumor tissue was mixed every 15 minutes to help digestion. After 2 h, 10 mL of HBSS with antibiotics were added and the cell suspension was strained over a 100 μm filter (Dutcher) which retained remaining tissue pieces. The suspension was centrifuged at 300 g for 5 minutes. In case of a visible red pellet, erythrocytes were lysed in 1 mL red blood cell lysis buffer (RBC, Invitrogen) for 5 min at room temperature. Then, the suspension was completed with 10 mL HBSS with antibiotics and centrifuged at 300 g for 5 minutes. The viable cell suspension was counted and 150,000 cells were used for the generation of tumoroids. The cells were resuspended in a reduced growth factor solubilized basement membrane matrix for Organoid Culture (Matrigel^®^, Corning) and plated as a drop in 24-well plates. The matrigel was allowed to solidify for 30 minutes in the incubator and then 500 μl of complete culture medium was added. The culture me-dium was composed of Advanced DMEM (Gibco) supplemented with 1X Glutamax, 10 mM Hepes, 1X Penicillin/Streptomycin, 1X Amphoteromicin, 50μg/mL Primocin, 1X B27 supplement, 5 mM Nicotinamide, 1.25 mM N-Acetylcystein, 250 ng/mL R-spondin 1, 5 nM Heregulinβ-1, 100 ng/mL Noggin, 20 ng/mL FGF-10, 5 ng/mL FGF-7, 5 ng/mL EGF, 500 nM A83-01, 500 nM SB202190 and 5μM Y-27632.

Tumoroids were split when confluent. Ice cold PBS was used to harvest tumoroids from the Matrigel. They were collected in a 15 mL falcon that was pre-coated with PBS containing 1 % BSA solution to prevent the tumoroids from adhering to the tube. The tumoroids were centrifuged at 300 g for 5 minutes and then digested with TrypLE solution (Gibco) for 5 min at 37 ° C. After enzymatic neutralization and washing, the tumoroid fragments were resuspended in Matrigel and reseeded as explained above to allow formation of new tumoroids.

Furthermore, after the initial digestion of the tumor tissue, 2 million cells were cryopreserved for the subsequent development of tumoroids. To create the tumoroids from the frozen cells, the vial was thawed slowly and the cells were centrifuged in 10 mL of HBSS with antibiotics at 300 g for 5 minutes. Then, the cells were counted and seeded in the same way as with fresh cells.

### 2.4. Freezing and thawing of tumoroids

Once the tumoroids were confluent, they were collected as described before, centrifuged at 300 g for 5 minutes, separated with a syringe mounted with a 21 G needle before being centrifuged again and frozen in 90% fetal bovine serum (FBS) and 10% DMSO.

Cryopreserved tumoroids were thawed slowly and 1mL of the thawing solution was added to the vial (Advanced DMEM (Gibco), 15 mM Hepes (Gibco), 1 % BSA (Sigma)). Then, the solution was transferred to a tube containing 2 mL of thawing solution. The tumoroids were centrifuged at 300 g for 5 minutes and the pellet of tumoroids was resus-pended with 30 μl of Matrigel and cultured as explained before.

### 2.5. HE, Immunohistochemistry and Immunofluorescence staining

The tumoroids were fixed in 2% paraformaldehyde with 0.1% glutaraldehyde for 24 h followed by dehydration in 20% sucrose for 24 h, embedding in gelatin and freezing at −80°C.

Standard H&E staining was carried out on 5 μm thick tumor and tumoroid sections to appreciate the cellular and tissue structure details, using Tissue-Tek Prisma^®^ Automated Slide Stainer. Images were acquired on a Nikon Eclipse NI-U with the Nikon Elements BR 4.50.00 software.

The immunohistochemical staining was carried out on 5 μm thick tumor and tumoroid sections using an automated protocol developed for the Discovery XT automated slide staining system (Ventana Medical Systems, Inc.). Tumor and tumoroid sections were in-cubated for 40 min with the appropriate antibody before incubation with Discovery UltraMap anti-Rabbit (760–4315, Roche) or anti-mouse horseradish peroxidase (HRP) (760–4313, Roche) secondary antibodies and the Discovery ChromoMap DAB kit reagents (760–159, Roche). Counterstaining and post-counterstaining were performed using hematoxylin and bluing reagent (Ventana, Roche Diagnostics). The following commercially available antibodies were used for the characterization: estrogen receptor (ER)–α (SC-8005, Santa Cruz), progesterone receptor (PR) (790–4296, Roche) and human epidermal growth factor 2 (HER-2) (790–4493, Roche).

The immunofluorescence staining was carried out on 12 μm thick tumor and tumoroid sections. Tumor and tumoroid sections were washed 3 times in PBS, pre-incubated in blocking buffer in 0.3% Triton, 5% Normal Donkey Serum (NDS) and 2% ovalbumin in PBS for 1 h at room temperature. Then the samples were incubated overnight at 4 °C with proliferation marker Ki67 (790–4286, Roche). After 3 washes with PBS, samples were incubated 1 h at 37 °C with secondary donkey anti-rabbit antibody conjugated to Alexa Fluor 488 (1:200, Invitrogen, Carlsbad CA, USA) in blocking buffer. They were rinsed with PBS and the cell nuclei were counterstained with Hoechst 33342 fluorescent dye (1/10000, Invitrogen, Carlsbad CA, USA) for 20 min at 4 °C. Finally, the tumor and tumoroid sections were mounted on the slide with Dako Fluorescent Mounting Medium (Agilent, Santa Clara CA, USA). Samples without the addition of primary antibody were used as negative control. The presented pictures are representative of independent triplicates.

### 2.6. Total protein extraction

Sections of fresh frozen tumor and corresponding tumoroids were collected in triplicate for each condition. The tumor sections and the tumoroids pellet were lysed with RIPA buffer (150 mM NaCl, 50 mM Tris, 5 mM EGTA, 2 mM EDTA, 100 mM NaF, 10 mM sodium pyrophosphate, 1% NP40, 1 mM PMSF, and 1X protease inhibitors) for total protein extraction. Three steps of 30 seconds sonication at amplitude 50% on ice was applied, cell debris were removed by centrifugation (16,000 × g, 10 min, 4°C), the supernatants were collected and protein concentrations were measured using a Bio-Rad Protein Assay Kit, according to the manufacturer’s instructions. To normalize the tumoroids and tumor protein quantities, 100 μg of each sample was used for protein digestion and subsequent shotgun proteomics analysis.

### 2.7. Shotgun proteomics

Protein digestion was performed using the FASP method(20). Briefly, reduction solution was added to the sample (100 mM DTT in 8 M urea in 0.1 M Tris / HCl, pH 8.5 (UA buffer)) and incubated for 15 minutes at 95°C. The protein solution was then loaded onto 10 kDa Amicon filters, supplemented with 200 μL of UA buffer and centrifuged for 30 min at 14,000 g. Next, 200 μL of UA buffer were loaded onto the filter and centrifuged for 30 min at 14,000 g. Then, 100 μL of alkylation solution (0.05 M iodoacetamide in UA buffer) were added and incubated for 20 min in the dark before centrifugation for 30 min at 14,000 g. Finally, a 50 mM ammonium bicarbonate solution (AB) was added and centrifuged again for 30 min at 14,000 g. This last step was repeated three time. For the digestion, 50 μL LysC/Trypsin at 20 μg/mL in AB buffer was added and incubated at 37°C overnight. The digested peptides were recovered after centrifugation for 30 min at 14,000 g. Then, two washes with 100 μL of AB buffer were performed by centrifugation for 30 min at 14,000 g. Finally, the eluted peptides were acidified with 10 μl of 0.1% trifluoroacetic acid (TFA) and dried under vacuum.

### 2.8. LC-MS/MS analysis

The samples once dried were reconstituted in 20 μL of a 0.1% TFA solution and desalted using a C18 ZipTip (Millipore, Saint-Quentin-en-Yvelines, France). After elution with 20 μL of 80 % acetonitrile (ACN)/ 0,1 % TFA, the peptides were vacuum dried. Samples were then reconstituted in 0.1 % formic acid/ACN (98:2, v/v), and separated by reverse phase liquid chromatography by an Easy-nLC 1000 nano-UPLC (Thermo Scientific) in the reverse phase using a preconcentration column (75 μm DI × 2 cm, 3 μm, Thermo Scientific) and an analytical column (Acclaim PepMap C18, 75 μm ID × 50 cm, 2 μm, Thermo Scientific) interfaced with a nanoelectrospray ion source on an Q-Exactive Orbitrap mass spectrometer (Thermo Scientific). Separation was performed using a linear gradient starting at 95 % solvent A (0.1% FA in water) and 5 % solvent B (0.1% FA in ACN) up to 70 % solvent A and 30 % solvent B for 120 min at 300 nL/min. The LC system was coupled onto a Thermo Scientific Q-Exactive mass spectrometer set to Top10 most intense precursors in data-dependent acquisition mode, with a voltage of 2.8 kV. The survey scans were set to a resolving power of 70 000 at FWHM (m/z 400), in positive mode and using a target AGC of 3E+6. For the shotgun proteomics, the instrument was set to perform MS/MS between +2 and +8 charge state.

### 2.9. Data analyses

All the MS data were processed with MaxQuant (version 1.5.6.5) software(21) using the Andromeda search engine(22). Proteins were identified by searching MS and MS/MS data against a database of Canis lupus familiaris obtain from Uniprot database and containing XXX sequences. For identification, the FDR at the peptide spectrum matches (PSMs) and protein level was set to 1%. Label-free quantification of proteins was performed using the MaxLFQ algorithm with the default parameters. Analysis of the proteins identified were performed using Perseus (version 1.5.6.0) software(23,24).

Multiple-sample tests were performed using ANOVA test with a p-value of 5% and preserving grouping in randomization. Visual heatmap representations of significant proteins variation were obtained using hierarchical clustering analysis. Functional annotation and characterization of identified proteins were obtained using PANTHER (version 13.0) software(25) and STRING (version 9.1)(26). The analysis of gene ontology, cellular components and biological processes, were performed with FunRich 3.0 analysis tool(27).

### 2.10. Tumoroid response to Paclitaxel

For tumoroid culture and drug response analysis, the same amount of tumoroids was dissociated with cold PBS. The pellet was then digested with TrypLE solution (Gibco) for 5 min at 37 ° C. The tumoroids were then diluted in HBSS and then passed through a 100 μm filter (Dutcher) to remove large tumoroids. Subsequently, the tumoroids were centrifuged at 300 g for 5 minutes and then suspended in 2% Matrigel/tumoroid culture medium (3-5000 tumoroids/mL). For the drug response, 100 μl of tumoroid solution was placed in wells of 96-well plates coated with 1.5 % agarose. The tumoroids were allowed to form during 72h and then treated with Paclitaxel for 7 days before performing the viability test. Cell viability was performed using CellTiter-Glo 3D (Promega) according to the manufacturer’s instructions and results were normalized to controls. Paclitaxel concentrations ranged from 0.01 μmol to 100 μmol (5 concentrations) and DMSO controls were added. After 7 days, 100 μL of CellTiter-Glo3D reagent (Promega, Madison, WI, USA) was added to each well and the plate was shaken at room temperature for 25 min. Luminescence was read on a TriStar2 S LB 942 Multimode Microplate Reader and the data were analyzed using GraphPad Prism 6.

## 3 Results

### 3.1. Feasibility of tumoroid culture from freshly resected canine mammary tumors

Canine mammary tumor was collected in the operating room at the time of tumor resection. For the characterization of the tumoroids’ cultures, 6 on 31 patients of the established biobank were included (**Supplementary Table 1**). For all of them, the resection was a primary mammary tumor. Patient’s age ranged between 5 and 14 years old and were all female. Based on the 2010 histologic classification for canine mammary tumors, the 6 tumors were annotated (**Supplementary Table 1**)(28). The 6 tumors were characterized with the most important and frequent biomarkers of breast cancer: estrogen receptor (ER), progesterone receptor (PR) and HER2. Among the 6 tumors, 4 have a triple-negative signature, signifying the absence of HER2, ER, and PR proteins expression (TM-02, TM-03, TM-05, TM-06) while 2 tumors are of luminal subtype with PR expression (TM-01) or PR/ER expression (TM-04) (**Supplementary Figure 1 and Supplementary Table 1**). In addition, the positive Ki67 labeling of each tumor was evaluated. We found that all 6 tumors showed Ki67 positive cells, with variable levels (**Supplementary Figure 2**).

After tumor resection, the tumor fragment was divided into three pieces: the first piece was kept fresh for tumoroid culture generation, the second one was frozen without prior fixation and the last piece was fixed in PFA and cryopreserved (**Figure 1A**). Frozen tissue was used for proteomics while fixed tissue was used for histology.

**Figure 1:**
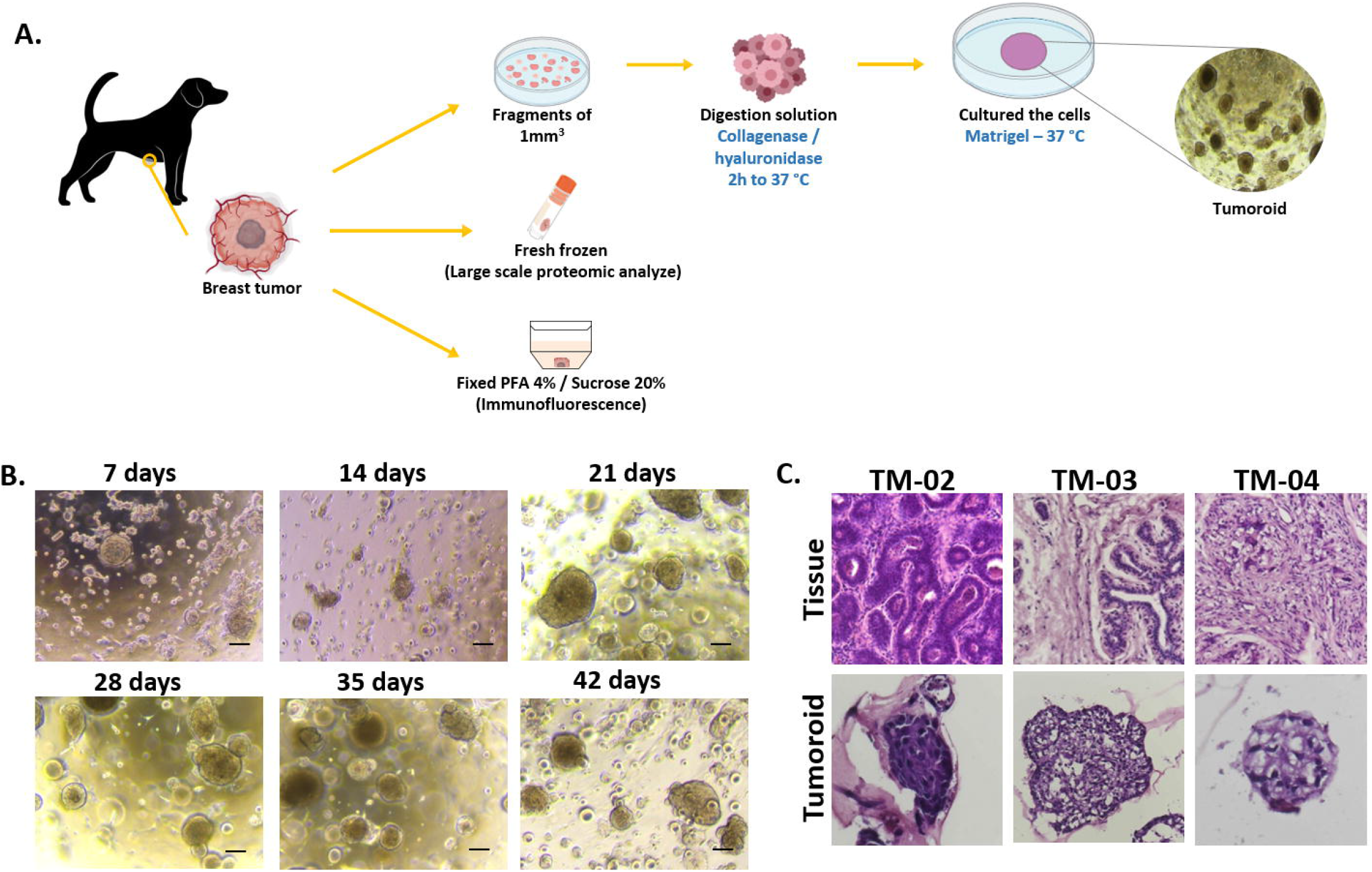
Breast cancer tumoroids culture established from canine patient. (A) Diagram presenting the strategy used to culture tumoroids from a canine mammary tumor. (B) Representative images of canine tumoroids at different time points of culture. Scale bar (200μm) is indicated. (C) H&E staining comparison between tumoroids and the tumor of origin for three different dog patients.

Fresh tissue pieces were mechanically and enzymatically dissociated to obtain single cell suspensions which were plated in Matrigel drops and overlaid with optimized mammary tumoroids culture medium. Cultures were followed by microscopy for evidence of tumoroids formation. We successfully generated tumoroid cultures from 31 of 33 tumor samples, an establishment success rate of 94%, with long-term expansion. Indeed, all tumoroids were grown for at least 42 days (6 passages) (**Figure 1B**). Majority of tumoroid lines were cryopreserved. The tumoroids morphologically reflected the original tumor they were derived from (**Figure 1C**). Tumoroids presented patient-specific heterogeneous morphologies, ranging from compact structures (TM-02) to more irregularly structures (TM-03 and TM-04).

### 3.2. Canine mammary tumoroids can be generated from both fresh and frozen cells and can be cryopreserved with similar histological and molecular features

Next, we wanted to evaluate whether tumoroids could be generated from frozen cells while keeping the same characteristics as fresh tumoroids. From the primary tumor sample, we divided the cell suspension into two parts: one part kept fresh for direct tumoroid formation, named “Fresh tumoroids” thereafter and the second part was frozen for indirect tumoroid formation, named “Tumoroids from frozen cells” thereafter. In addition, in order to characterize our biobank, we wanted to make sure that cryopreservation did not affect the tumoroids features. We therefore compared these two types of tumoroid cultures to thawed tumoroids, named “Frozen tumoroids” thereafter (**Figure 2A**). Tumoroids from these different culture conditions were left in culture during 4-5 weeks (date 1) or 6-7 weeks (date 2) and compared to study tumoroids drift overtime.

**Figure 2:**
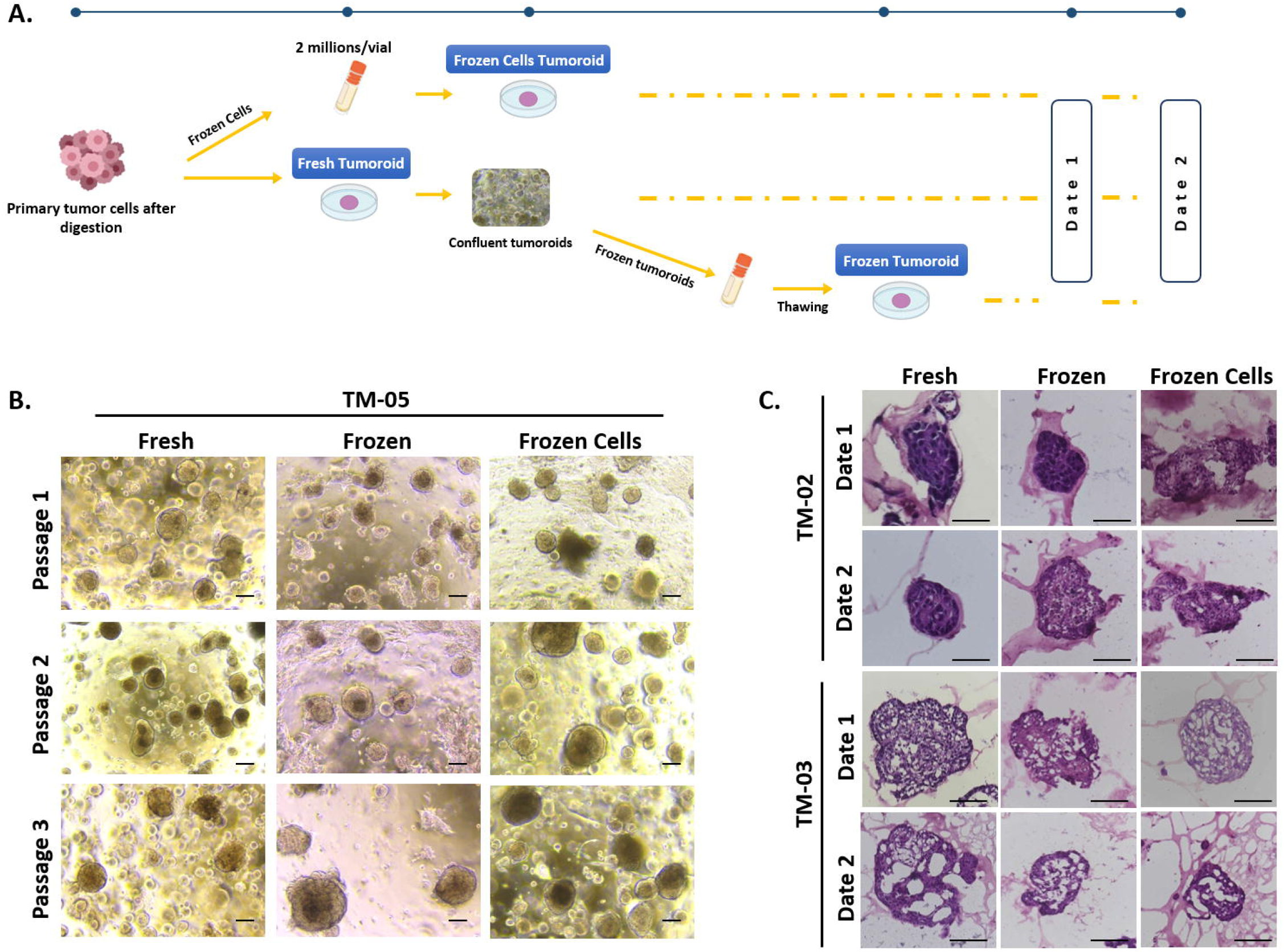
Study of canine mammary tumoroids drift. (A) Diagram showing the strategy used to generate the different types of tumoroids. After tumor digestion, a part of the cells were frozen and then thawed to generate the “Frozen Cell Tumoroids”. The remaining cells were used to generate the “Fresh tumoroids”. A part of these tumoroids were cryopreserved and then thawed, corresponding to the “Frozen Tumoroids”. The three types of tumoroids were compared at the same time point post-culture at date 1 (4-5 weeks) or date 2 (6-7 weeks). (B) Representative images of the three types of canine tumoroids at passage one, two and three after Date 1. Scale bar (200μm) is indicated. (C) H&E staining comparison of fresh, frozen and frozen cells tumoroids. Scale bar=100 μm

Histologic and molecular drifts of tumoroids after cryopreservation and after long-term culture were studied. First, the culture of tumoroids was successful for each culture condition and after serial passages as well (**Figure 2B**). Tumoroid formation efficiency was not found to be different between cryopreserved cells and fresh cells. Tumoroid cultures from fresh and frozen cells could be similarly long-term cultured and passaged (**Figure 2B**). At the histological level, tumoroids derived from fresh cells, frozen cells or after cryopreservation retained the same architecture. **Figure 2C** presents representative images of H&E staining of tumoroids derived from two different tumors. Tumoroids derived from TM-02 were compact while tumoroids derived from TM-03 were more irregular whatever the culture condition and time in culture. The freezing procedure did not affect tumoroids morphology.

The ER, PR and HER2 expression profiles of breast cancer tumoroids were compared with their original breast cancer tissues. For this, 2 tumors were used: TM-03 (triple negative subtype) and TM-04 (luminal subtype) (**Figure 3**). The results showed that the tumoroids maintain the same expression profile of the tumor of origin. In the case of TM-03, tumoroids present a triple negative subtype as the tumor they are derived from. In the case of TM-04 tumoroids, we can observe ER and PR positive cells similar to the tumor of origin.

**Figure 3:**
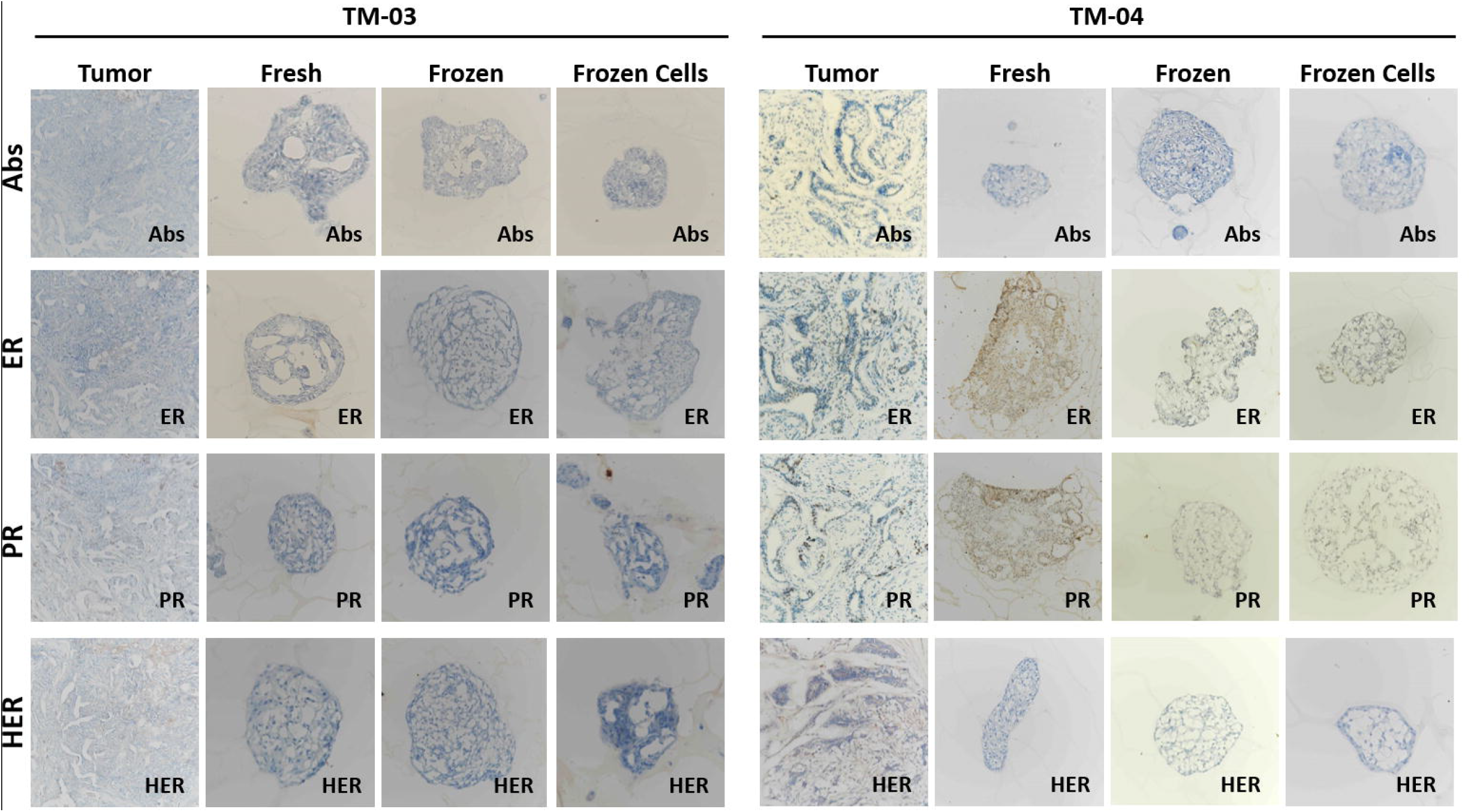
Histology and receptor status (ER, PR, HER2) of breast cancer tumoroids. Comparative histological and immunohistochemical images of breast cancer tumoroids and their original breast cancer tissues.

Finally we verified if the proliferation of the tumoroids in all three conditions was similar. We used the TM-05 tumoroids that show a large number of positive cells in the tumor of origin (**Supplementary Figure 2**) to answer this question. The proliferative activity of the tumoroids was determined by the percentage of Ki67+ cells with respect to the total cells of each tumoroids conditions. Proliferation activity of tumoroids do not show a significant difference between Fresh (8.61%), Frozen (6.96%) and FrozenCell (6.29%) tumoroids (**Figure 4**).

**Figure 4:**
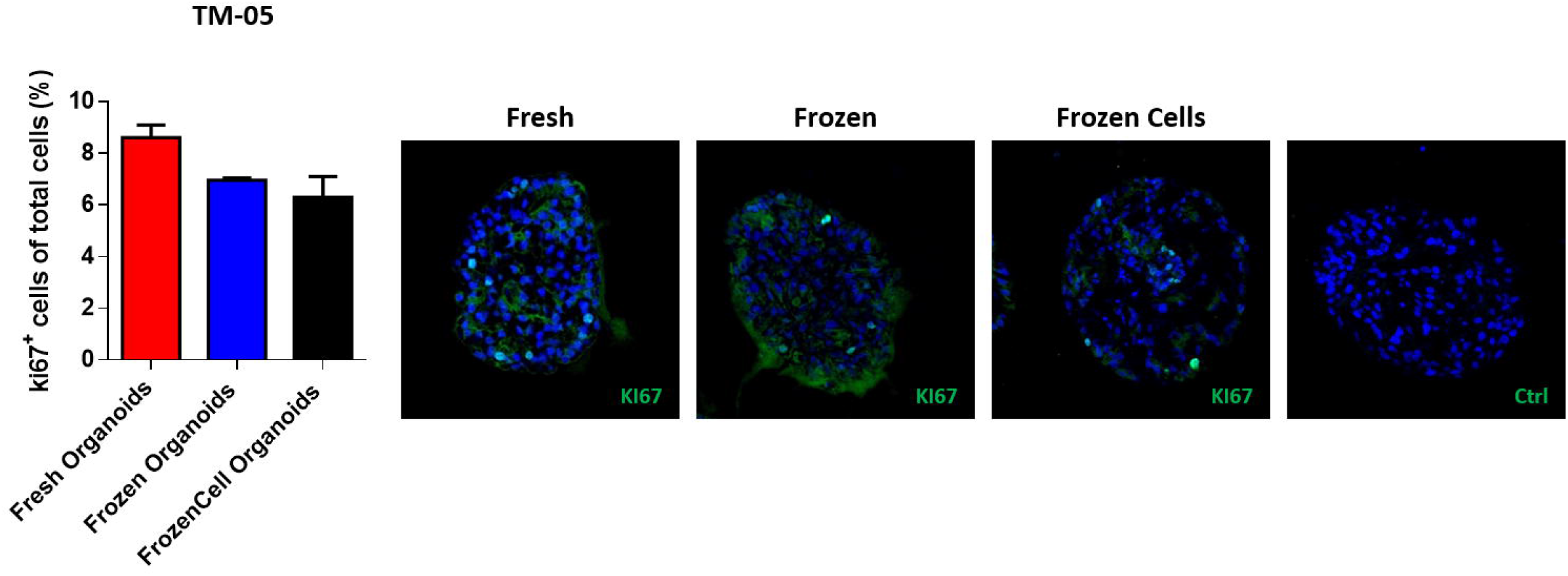
Proliferative activity of the tumoroids. Comparative quantification of the percentage of Ki67+ cells in tumoroids (Fresh, Frozen and FrozenCell conditions). The average of the triplicates is shown and error bars mean SD. Proliferation did not differ significantly between the 3 conditions.

### 3.3. Similar proteomic profiles are observed between tumoroids generated from fresh and frozen cells over time while cryopreservation seems to trigger a more pronounced molecular drift

We have shown that the freezing procedure as well as the passages did not impact the morphology of tumoroids neither their histological features. In order to understand, whether the frozen tumoroids or tumoroids made from frozen cells kept similar molecular profiles as the fresh tumoroids or the original tumor, we have performed a large-scale unbiased proteomic analysis. The study was carried out on 3 tumors: TM-01, TM-02 and TM-03; of which the three types of tumoroids were made and compared with each other, and with the original tumor. For this the extracted proteins were quantified and the same amount of proteins was used. In addition, to understand if there was any molecular drift over time, we have analyzed the proteome of tumoroids at two different dates. More than 2,500 proteins were identified in total through biological replications within the experimental groups.

First, taking into account the two dates of tumoroid passage (D1 and D2), 1796 proteins were identified shared by the six conditions; D1-Fresh, D1-FrozenCell, D1-Frozen, D2-Fresh, D2-FrozenCell and D2-Frozen (62% of all the proteins identified) (**Figure 5A**) (**Supplementary Table 2**). The D1-Frozen tumoroids seem to be the most different compared to all the other conditions as shown on **Figure 5A**. In fact, 343 proteins were identified in all conditions except in D1-Frozen. This may be due to a lack of protein diversity in this condition, as all samples were quantified to have the same amount of proteins. However, if we observed more closely the proteins lacking in the condition of D1-Frozen (**Figure 5B**), we found several proteins involved in metabolism and energy pathways such as HMGCS2 (Hydroxymethylglutaryl-CoA synthase), SOAT1 (Sterol O-acyltransferase 1), PRPSAP1 (Phosphoribosyl pyrophosphate synthase-associated protein 1), ECI1 (Enoyl-CoA delta isomerase 1), MMP9 (Matrix metalloproteinase-9), UBE4B (Ubiquitin conjugation factor E4 B), BMP1 (Bone morphogenetic protein 1) and HEXB (Beta-hexosaminidase subunit beta) among many others. In addition, we found proteins involved in the inhibition of apoptosis such as API5 (Apoptosis Inhibitor 5), SOD2 (Superoxide dismutase 2), SYVN1 (Synoviolin 1). We also identified a lot of proteins involved in Cell growth and/or maintenance and Cell communication. Very interestingly, proteins involved in immune response processes were enriched such as NRP1 (Neuropilin 1), PROCR (Protein C Receptor-CD201), ALCAM (activated leukocyte cell adhesion molecule-CD166), CD109 (Cluster of Differentiation 109), LBP (Lipopolysaccharide-binding protein), ST6GAL1 (ST6 Beta-Galactoside Alpha-2,6-Sialyltransferase 1), LRRC8A (Leucine-rich repeat-containing protein 8A) and CFB (Complement Factor B). These proteins which were not found in the D1-Frozen condition have a proliferative, immune and anti-apoptosis profile; demonstrating a lack of these biological processes in the D1-Frozen condition.

**Figure 5:**
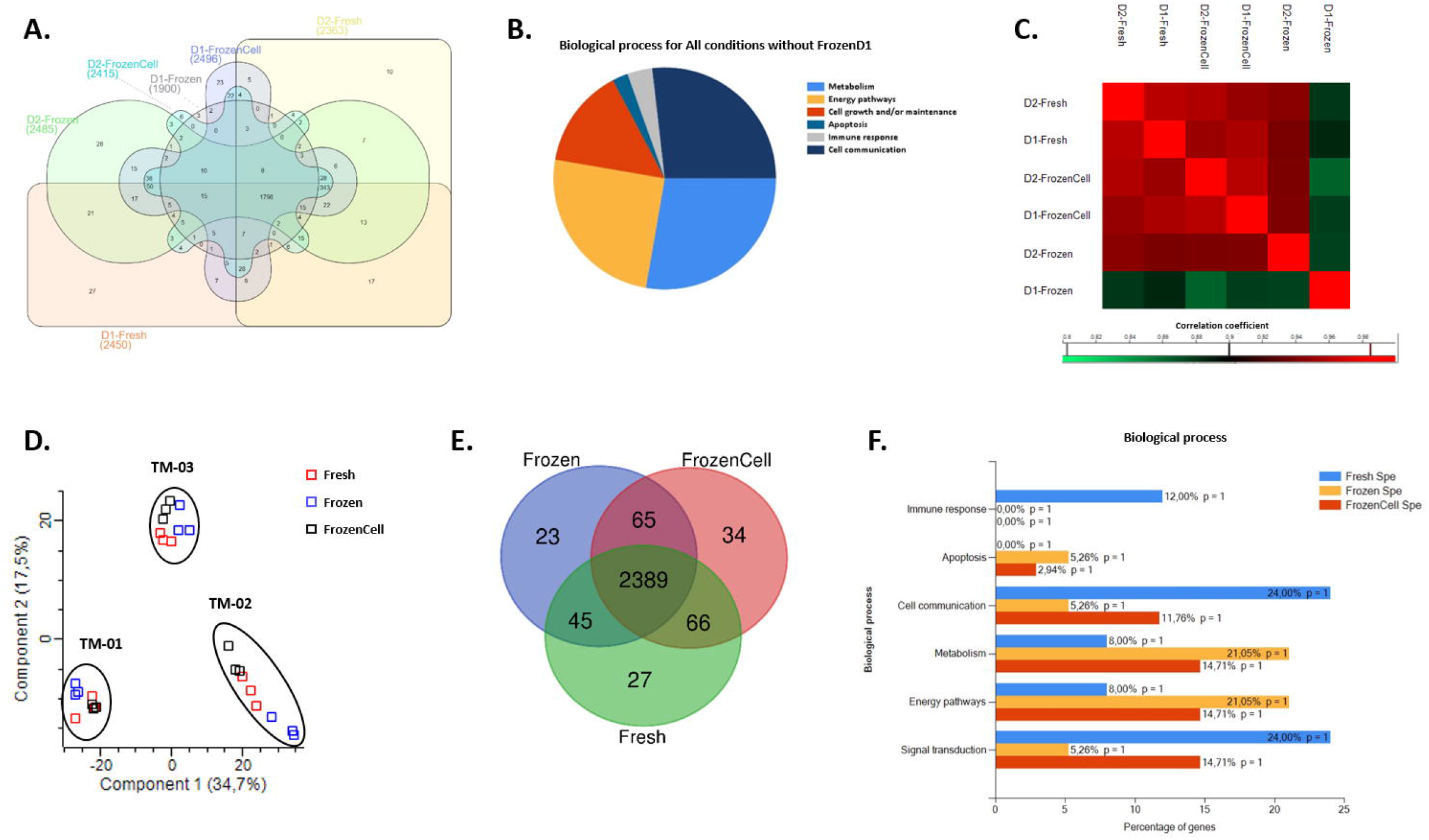
Proteomics analysis of CMT tumoroids. (A) Venn diagram representing specific proteins identified in the Fresh, Frozen and FrozenCell tumoroids at Date 1 and Date 2. (B) Biological processes of the proteins identified in common except in Frozen D1 tumoroids. (C) Matrix correlation studies between the two dates in the tree different conditions. (D) PCA analysis of the proteomics data from the tree different tumoroid conditions. (E) Venn diagram representing specific proteins in the Fresh, Frozen and FrozenCell tumoroids. (F) Funrich biological process distribution of the specific proteins identified in Fresh, Frozen and FrozenCell tumoroids.

Likewise by a Pearson correlation analysis, hierarchical clustering of all the samples based on the correlation coefficients between them revealed higher similarity between Fresh and Frozen Cell tumoroids at date 1 and 2. Frozen tumoroids were more different, specifically at date 1 **(Figure 5C)**. The similarity of D1-Frozen with the other conditions was less than 87% while all the other conditions showed more than 95% similarity. The duration of the tumoroids culture did not seem to have a big impact on their proteomic profiles. The fact that D1-Frozen tumoroids were more distinct suggests that the tumoroids should be preferentially left in culture long enough to recover after freezing, which was not observed from D1-FrozenCells.

Knowing that the time in culture did not impact too much their molecular profile, we then wanted to verify whether the culture condition impacted or not their proteome. For that, we have compared the proteomic profiles of tumoroids from three culture conditions: fresh, frozen and tumoroids made from frozen cells. First of all, the principal component analysis (PCA) based on the LFQ values of the protein identification showed that the samples were grouped by tumor and not according to the type of culture condition (**Figure 5D**). This sample grouping by PCA means that there was a high level of similarity between the biological replicates of each condition but also between the tumoroids without influence of their culture condition. Furthermore, a Venn diagram showing the number of common and unique proteins in all conditions showed that a majority of proteins were identified in all three conditions of culture (2389 proteins, representing 90% of all proteins). However, some proteins were found specifically expressed in each condition: 27 identified specifically in fresh tumoroids, 23 in frozen tumoroids and 34 in tumoroids made from frozen Cells (**Figure 5E**) (**Supplementary Table 3**). Based on the GO terms enrichment analysis of the biological processes using FunRich software, we observed that these proteins, specifically expressed in each condition, were linked to different biological processes (**Figure 5F**). An enrichment of proteins linked to metabolic and energy signaling pathways was found in Frozen and FrozenCell tumoroid conditions compared to Fresh tumoroids, such as AMY1A (amylase, alpha 1A), SDR9C7 (short chain dehydrogenase/reductase family 9C-member 7), CDA (cytidine deaminase) and ARG1 (arginase 1), FKBP (FK506 binding protein), NDUFB10 (NADH dehydrogenase (ubiquinone) 1 beta subcomplex), DDO (D-aspartate oxidase), ADH5 (alcohol dehydrogenase 5 (class III)) and SIAE (sialic acid acetylesterase). In addition, in the Frozen and FrozenCell condition, we have identified proteins involved in apoptosis like the apoptosis facilitator BCL2L13 (Bcl-2-like protein 13), ATG5 (autophagy related 5) and TXNRD2 (thioredoxin reductase 2). In the Fresh tumoroids, a higher number of proteins linked to cell communication and to signal transduction were identified. Interestingly, some of the specific proteins identified in fresh tumoroids were involved in the immune response, such as GZMB (Granzyme B) expressed by cytotoxic T and NK cells, the cell adhesion molecule Siglec1 (Sialoadhesin) expressed by macrophages, as well as CD163 (Cluster Differentiation 163), a marker of anti-inflammatory macrophages and the AMBP (alpha-1 microglobulin/bikunin) precursor of a glycoprotein synthesized by lymphocytes. CD177 (CD177 molecule), a marker of neutrophil activation, was also identified specifically in fresh tumoroids.

In order to better understand the differences linked to the culture conditions, an analysis of the variation of abundance of common proteins to all conditions (2389 proteins) was carried out, using a multiple sample test ANOVA with an FDR of 0.05. A total of 489 proteins showed significantly different expression between the three groups. These specific variations were analyzed by hierarchical clustering and then illustrated by a Heatmap (**Figure 6A**). Six clusters of proteins were identified: one cluster representing the specific underexpressed proteins and one representing the specific overexpressed proteins for each condition (**Supplementary Table 4**). Based on over- and under-expressed proteins, fresh tumoroids and tumoroids made from frozen cells showed more similarities compared to frozen tumoroids, as observed before as well. In order to understand more precisely the impact of these proteins, the analysis of the GOterms of each cluster was carried out with Cytoscape and ClueGO software, allowing to generate the networks connecting the proteins overexpressed (in red) and underexpressed (in green) to their biological processes.

**Figure 6:**
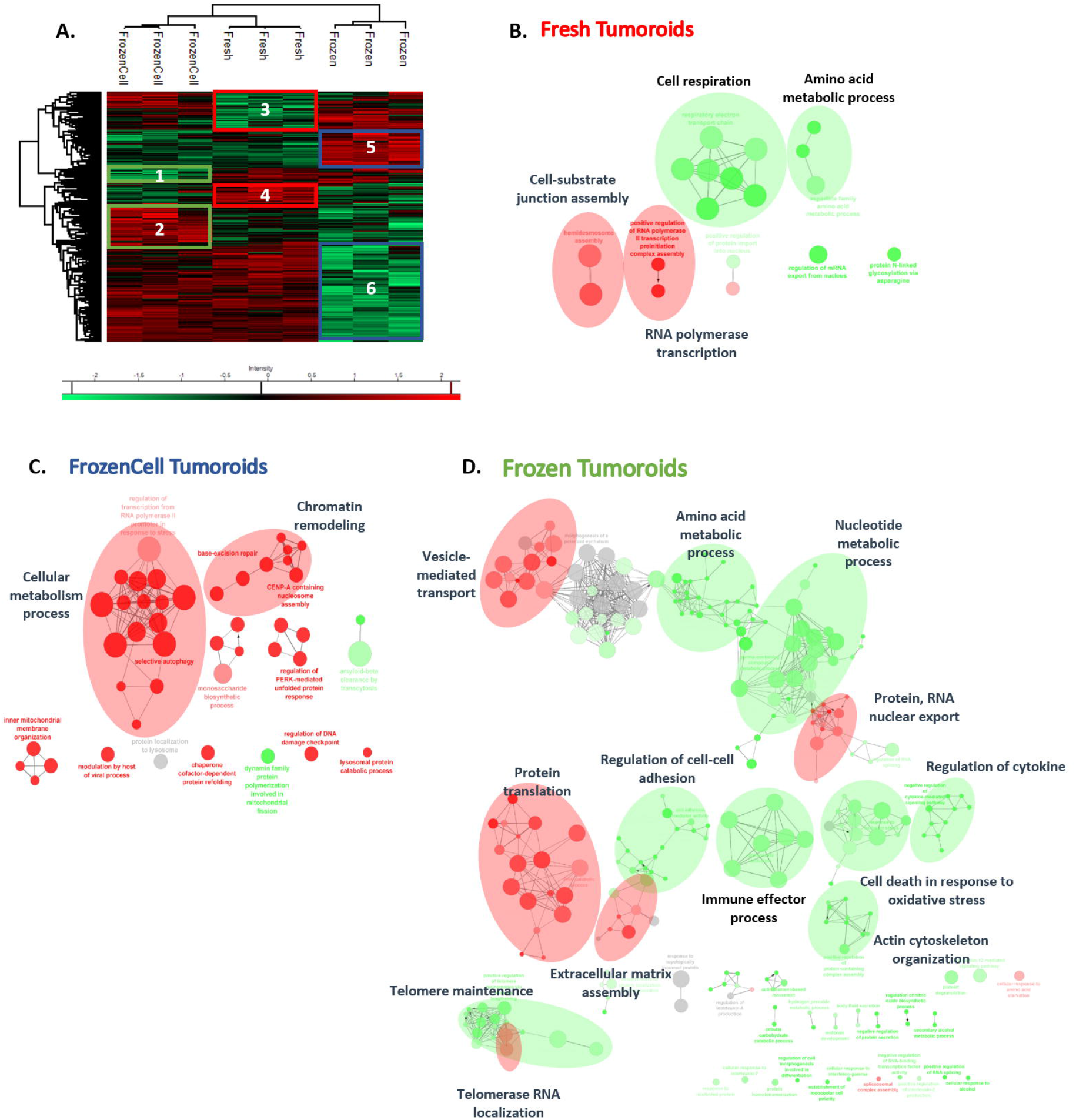
Proteomics analysis of Fresh, Frozen and FrozenCells tumoroids. (A) Hierarchical clustering of the most variable proteins between the 3 conditions (n = 3 for each condition, ANOVA with permutation-based FDR < 0.05). Network of proteins overexpressed (red) or underexpressed (green) in Fresh (B), FrozenCell (C) and Frozen (D) tumoroids and their associated GO terms. The networks were enriched through addition of STRING network to the identified proteins using ClueGO application on Cytoscape.

The results showed that in the fresh condition (Clusters 3 - 4) (**Figures 6A and 6B**), the signaling pathways linked to cellular respiration and to amino acid metabolism were underexpressed while the frozen tumoroids and FrozenCell tumoroids express more proteins in these two biological processes, which can be explained by the cryopreservation. In another hand, the assembly of the cell-substrate junction and the RNA translation by RNA polymerase appeared to be overexpressed in the Fresh condition compared to the other conditions, involving proteins like ITGB4 (Integrin beta-4), In order to better understand the differences linked to the culture conditions, an analysis of the variation of abundance of common proteins to all conditions (2389 proteins) was carried out, using a multiple sample test ANOVA with an FDR of 0.05. A total of 489 proteins showed significantly different expression between the three groups. These specific variations were analyzed by hierarchical clustering and then illustrated by a Heatmap (**Figure 6A**). Six clusters of proteins were identified: one cluster representing the specific underexpressed proteins and one representing the specific overexpressed proteins for each condition. Based on over- and under-expressed proteins, fresh tumoroids and tumoroids made from frozen cells showed more similarities compared to frozen tumoroids, as observed before as well. In order to understand more precisely the impact of these proteins, the analysis of the GOterms of each cluster was carried out with Cytoscape and ClueGO software, allowing to generate the networks connecting the proteins overexpressed (in red) and underexpressed (in green) to their biological processes.

The results showed that in the fresh condition (Clusters 3 - 4) (**Figures 6A and 6B**), the signaling pathways linked to cellular respiration and to amino acid metabolism were underexpressed while the frozen tumoroids and FrozenCell tumoroids express more proteins in these two biological processes, which can be explained by the cryopreservation. In another hand, the assembly of the cell-substrate junction and the RNA translation by RNA polymerase appeared to be overexpressed in the Fresh condition compared to the other conditions, involving proteins like ITGB4 (Integrin beta-4), Macf1 (Microtubule-actin cross-linking factor 1), PSMC2-6 (proteins linking with proteasome), KRT14 (keratin 14), PLEC (plectin) and VCL (Vinculin). The proteins involved in cell adhesion are overexpressed in the Fresh condition, which can explained by the formation of tumoroids that form their own extracellular matrix and by the cell compaction.

For the FrozenCell condition (Clusters 1 - 2) (**Figure 6C**), there is a higher abundance of proteins linked to chromatin remodeling and cellular metabolism as we observed before, some examples of proteins are UBA52 (Ubiquitin-60S ribosomal protein L40) and PSMC1, PSMD5 (26S proteasome non-ATPase regulatory subunit 1-5) proteasome regulatory forms, RAB7A (Ras-related protein Rab-7a), HSPA9- HSPA5- HSPA8 (Endoplasmic reticulum chaperone BiP), DDB1 (DNA damage-binding protein 1), MDH2 (Malate dehydrogenase), SLC25A12 (Calcium-binding mitochondrial carrier protein Aralar1). Again, a high metabolic activity that is a consequence of freezing, in addition to the chromatic remodeling that is involved in the cell division cycle, can be linked to a process of multiplication and recovery from freezing that seems important in this condition.

Finally, in frozen tumoroids (Clusters 5 - 6) (**Figure 6D**), many proteins related to protein translation, the proteins of the Extracellular matrix assembly, the Vesicle-mediated transport, Protein translation and Protein - RNA nuclear export were found to be overexpressed. We find an overexpression of proteins linked to the transport of extracellular vesicles, vesicle budding from membrane, vesicle targeting, vesicle coating and COPPII coated vesicles cargo such as: ARCN1 (Coatomer subunit beta-delta), AP2A1 (AP-2 complex subunit alpha), DYNC1H1 (Cytoplasmic dynein 1 heavy chain 1), AP2B1 (AP complex subunit beta), ANXA7 (Annexin A7), SEC13 (Protein SEC13 homolog), among others. In addition we can observe an overexpression of the biological processes linked to Protein translation, Protein-RNA nuclear export and Telomerase RNA localization. For example, different proteins of Eukaryotic translation initiation factor (4A-III, 3 subunit A, 3 subunit L, 3 subunit B, 3 subunit E, among others) and 40S and 60S ribosomal protein (RPL10, RPL13A, RPL14, RPL15, RPS11, RPS13, RPS18, RPS28, RPS3, RPS8, RPSA) are overexpressed in the Protein translation biological process. These biological processes show a dysfunction in the translation pathways that we know contribute to cancer progression, for example, in the deregulation of ncRNAs that leads to aberrant protein translation in cancers(29).

On the contrary, the underexpressed proteins are related to the metabolism of amino acids or nucleotides, negative regulation of cytokines, immune effector process, and the organization of the cytoskeleton, cell adhesion and death. Different metabolic pathways were touched, such as dicarboxylic acid metabolic process, purine ribonucleotide biosynthetic process, pyruvate metabolic process, generation of precursor metabolites and energy. Regarding the organization of the cytoskeleton and cell adhesion, different isoforms of laminin, collagen, catenin and Coronin-1B were found to be under expressed in this condition. Apoptosis and cell death proteins were also found under expressed as CYP1B1 (Cytochrome P450), HSPA1 (Heat shock protein 75 kDa), ARL6IP5 (PRA1 family protein 3), TRAP1 (TNF receptor associated protein 1), among others. In addition, we have identified underexpressed proteins linked to a regulation of cytokines and to the immune effector process, in which we find proteins such as: CD44 (CD44 antigen), thrombospondin-1, SAMHD1 (Deoxynucleoside triphosphate triphosphohydrolase SAMHD1), TINAGL1 (Tubulointerstitial nephritis antigen like 1), GAA (Alpha glucosidase), LGALS9 (Galectin), among others.

The Cytoscape and ClueGO analysis shows that Fresh and FrozenCell conditions have limited amount of underexpressed and overexpressed proteins, while frozen condition shows three time more proteins with significant variation. On the other side, even if the FrozenCell is closer to Fresh Tumoroids than Frozen condition, the degree of similarity stays high.

### 3.4. The proteome of canine mammary tumoroids is very similar to the tumor they originate and therefore represent a faithful breast cancer model

We next wanted to determine whether the three different types of tumoroids were similar to the tumor of origin, since the tumoroids will be used as a model of breast cancer.

For this, a Venn diagram (**Figure 7A**) was made and showed the number of common and unique proteins in all conditions (**Supplementary Table 5**). It can be observed that a majority of proteins were identified in all three conditions of culture (2138 proteins, representing 74% of all proteins). However, there were specific proteins for each condition especially in the original tumor: 4 identified specifically in fresh tumoroids, 15 in frozen tumoroids, 15 in tumoroids made from frozen cells and 153 specific proteins that were found specifically in the tumor. These 153 proteins are involved in different biological processes such as Cell growth and/or maintenance, Cell communication, Signal transduction and Immune response (**Figure 7B**). Interestingly, an immunological profile can be observed in the tumor compared to the tumoroids. We found many proteins involved in the complement signaling pathway (complement factor I, complement component 4 binding protein, complement component 5, complement component 7 and complement component 8) that are involved in immunological response and phagocytosis and found overexpressed in different types of cancer, such as breast cancer(30). In addition, proteins such as CD93 molecule, CD34 molecule, C-type lectin domain family 4 member G, haptoglobin and joining chain of multimeric IgA and IgM have been identified and are all implicated in immune response. The AOC3 (amine oxidase, copper containing 3) protein was also identified, and has been recently described to play a role in the reduction of immune cell recruitment and impacting the promotion and progression of lung cancer(31).

**Figure 7:**
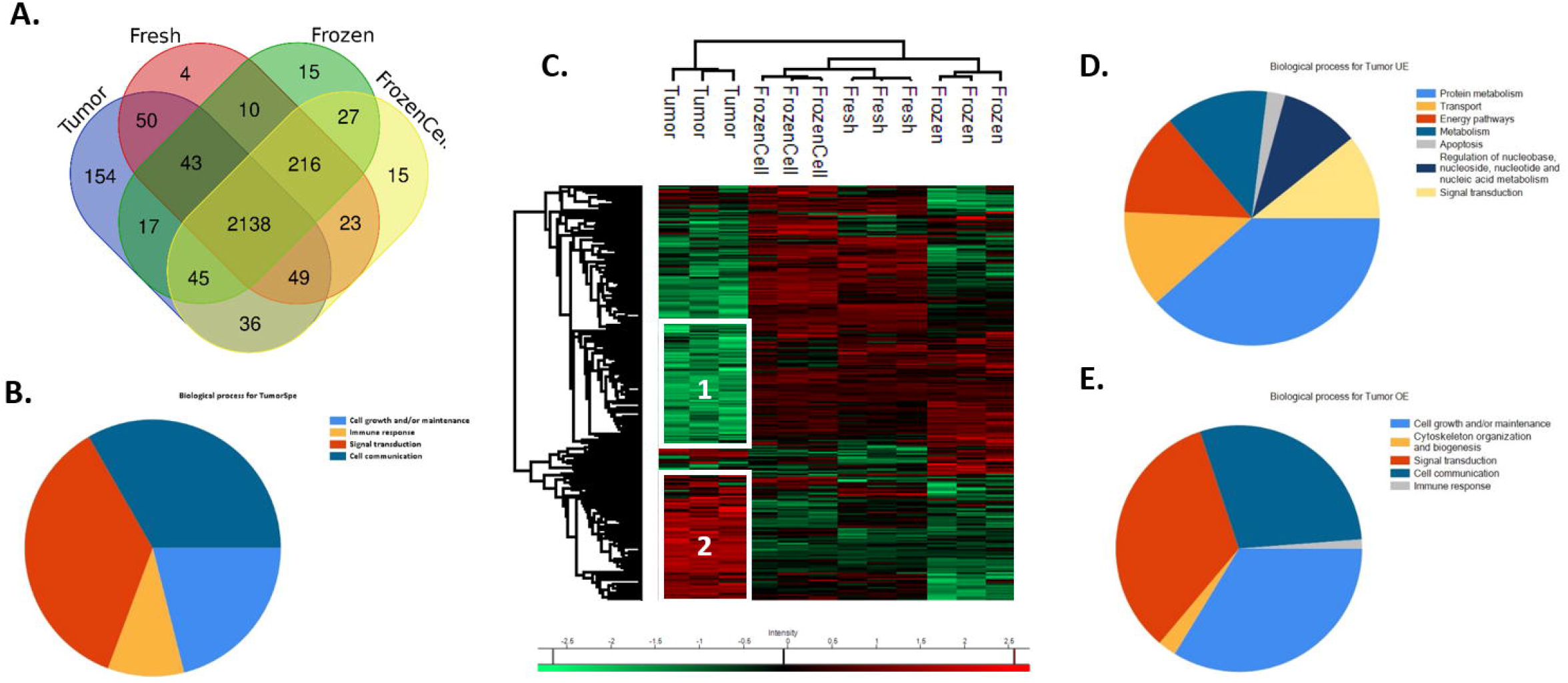
Proteomics analysis comparing the primary tumor to their derived tumoroids. (A) Venn diagram representing specific proteins in tumor of origin, Fresh, Frozen and FrozenCell tumoroids. (B) Biological processes of the specific proteins identified in primary tumors using Funrich and ClueGO. (C) Hierarchical clustering of the most variable proteins between the tumor of origin and the 3 tumoroid conditions (n = 3 for each condition, ANOVA with permutation-based FDR < 0.05). Biological processes distribution of underexpressed (D) and overexpressed (E) proteins in tumors compared to tumoroids using Funrich and Cluego.

To better understand the differences between tumor and tumoroids, an analysis of the variation of abundance of common proteins was carried out, using a multiple sample test ANOVA with an FDR of 0.05. A total of 512 proteins showed significantly different expression between the four groups. These specific variations were analyzed by hierarchical clustering and then illustrated by a Heatmap (**Figure 7C**). The HeatMap shows only small variations between the three types of tumoroids. Frozen tumoroids were more different compared to the two other culture conditions confirming the previous results. Interestingly, a small cluster of overexpressed proteins was observed in Tumor, Fresh and FrozenCell tumoroids, while down-expressed in Frozen tumoroids. This result shows again that the conditions more similar to the tumor of origin are the Fresh and FrozenCell tumoroids.

We next focused on the two clusters that showed the significant differences between the tumor of origin and the tumoroids (**Supplementary Table 6**). The Heat Map shows two clusters of proteins over- or down-expressed in Tumor compared to the different tumoroids. Functional annotation and characterization of these two clusters of proteins were performed using FunRich software. The results showed that the biological processes underexpressed in the tumor compared to tumoroids are different processes involved in metabolism, transport, energy pathways, apoptosis and signal transduction (**Figure 7D**). On the contrary, we can observe that proteins overexpressed in the tumor compared to tumoroids are involved in cell growth and maintenance, cytoskeleton organization, cell communication, signal transduction and immune response (**Figure 7E**). These results confirm that in tumoroids we find a higher metabolic activity, especially in frozen tumoroids as demonstrated above. In addition, we can observe apoptotic proteins such as BCL2-associated athanogene 6, heat shock 60kDa protein 1, PH domain and leucine rich repeat protein phosphatase 2, cell cycle and apoptosis regulator 2, underexpressed in the tumor and consequently overexpressed in the tumoroids. In addition, in the tumor we can find an important proliferative profile demonstrated by the overexpression of proteins involved in cell communication and linked to the organization of the cytoskeleton and cell growth (actin beta-like 2, collagen type VI, tubulin, beta 4B class IVb, lamin A/C, actin alpha 2, actin related protein, among others). Finally, we can find an immune profile more present in the tumor of origin.

### 3.5 Canine mammary tumoroids can be used to test human drugs and cryopreservation of tumoroids does not impact drug response

In order to evaluate canine mammary tumoroids as useful tools for translational *in vitro* drug screening studies, we performed cell viability assays in presence of a chemotherapeutic agent used in human medicine, Paclitaxel. Tumoroids were treated with Paclitaxel for 7 days before cell viability was measured. Representative images of tumoroids derived from TM-05 tumor are shown in **Figure 8A** demonstrating drug sensitivity depending on the drug concentration. Using 6 concentrations of Paclitaxel, we generated dose-response curves (**Figure 8B**). First, we could demonstrate that tumoroids derived from fresh canine mammary tumors responded well to Paclitaxel with an IC50 ranging from 0.1 and 1 μM. 0.1 μM Paclitaxel was needed to kill 50% of tumor cells for TM-04 and TM-06 while around 1 μM was needed for TM-05 demonstrating higher resistance to Paclitaxel.

**Figure 8:**
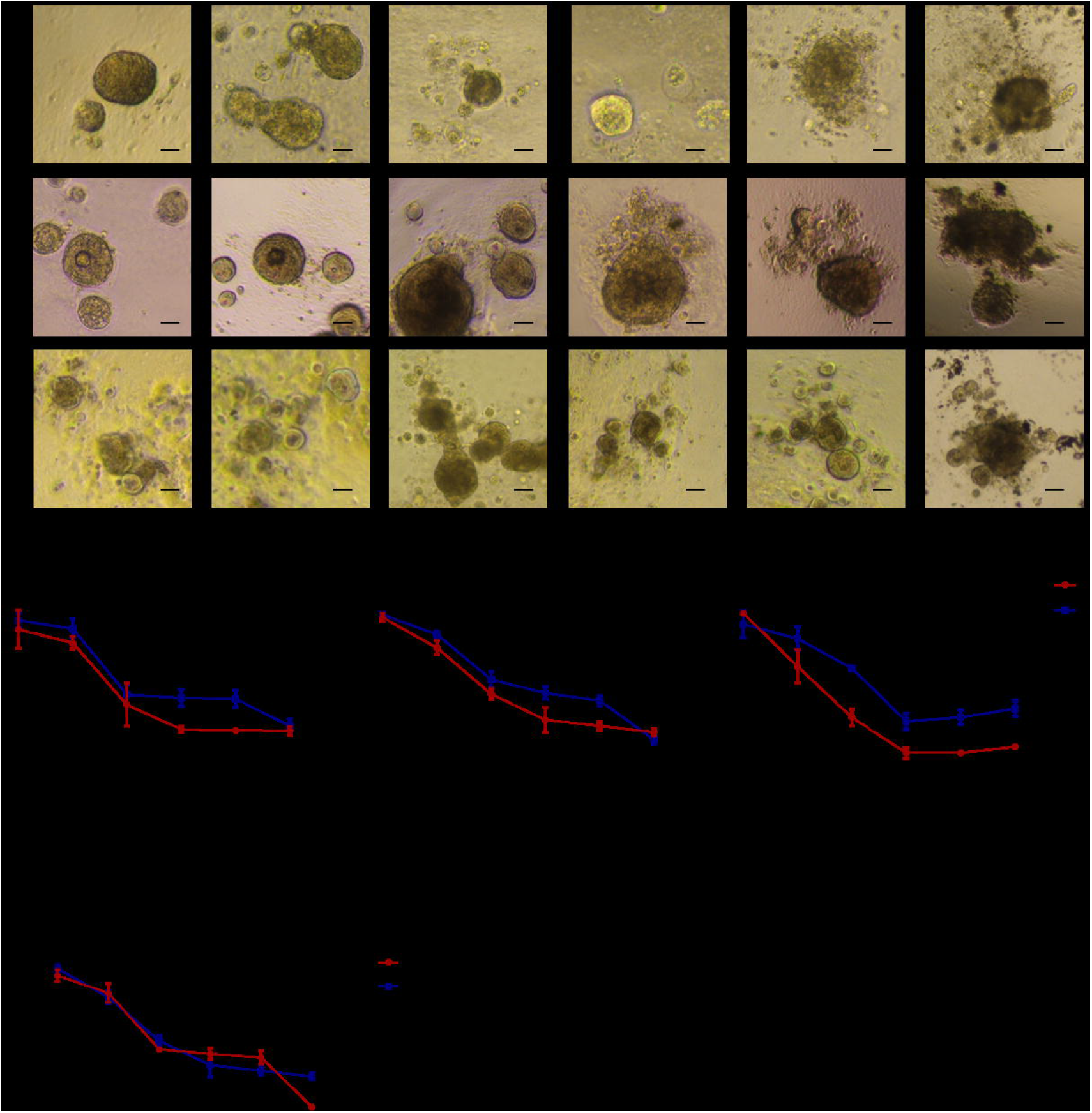
Drug response of canine and human tumoroids to Paclitaxel. (A) Representative bright field images showing the morphology of the three types of tumoroids after 7 days treatment with Paclitaxel at different concentrations. Scale bar (100μm) is indicated. (B) Quantification of the tumoroids viability following Paclitaxel treatment. Tumoroids were generated from three different canine mammary tumors (TM-04, TM-05 and TM-06) and drug response was compared between fresh, frozen and frozen cell tumoroids. (C) Tumoroids were generated from human and canine mammary tumors and drug response was compared between human and canine tumoroids. Different concentrations of drug were used and compared to non-treated tumoroids. Data are the means ± SD.

We next compared the Paclitaxel response between fresh tumoroids, frozen tumoroids and tumoroids made from frozen cells in order to verify whether cryopreserved tumoroids could represent faithful models for drug testing. Killed curves from these three culture conditions were similar for the three tumors tested (**Figure 8B**). As we have observed before with the proteomic analysis, fresh tumoroids and tumoroids derived from frozen cells were the most similar in term of Paclitaxel responses. Nonetheless, tumoroids derived from frozen cells appear to become slightly more resistant at higher concentrations of Paclitaxel (**Figure 8A and B**) compared to fresh tumoroids. Indeed, an increase of viability of tumoroids derived from frozen cells can be observed for each tumor at a concentration of 100 μM. A 50% viability of tumoroids was measured in this condition while only 35% of cells were viable in fresh tumoroids (**Figure 8B**). Finally, the response to Paclitaxel of frozen tumoroids appears to be slightly different compared to the two other conditions, even if not significant. For TM-04 and TM-06, fresh tumoroids and tumoroids made from frozen cells seem to be more sensitive whatever the concentration of paclitaxel used compared to frozen tumoroids. For TM-05, the three curves are more similar. A 50% viability decrease of fresh tumoroids and tumoroids made from frozen cells is observed between 0.1 to 1 μM of drug while between 1 and 10 μM of Paclitaxel are needed to kill 50% of tumor cells in the frozen tumoroids condition (**Figure 8B**). In the end, we have shown that Paclitaxel response between luminal subtype CMT tumoroids and human breast tumoroids was similar with 0.1μM of Paclitaxel needed to kill 50% of tumor cells (**Figure 8C**).

In conclusion, canine tumoroids respond well to a chemotherapeutic agent used in human medicine. The way the tumoroids are preserved has little impact on drug response. It seems, however, that tumoroids made from frozen cells best mimic the drug response of fresh tumoroids.

## 4 Discussion

Tumoroids provide an alternative to pre-clinical animal experiments and can help predict tumor response to therapy and screen new drugs. Until now, breast cancer tumoroids have been derived mainly from murine and human tissues(3,32). However, murine tumors do not reliably reflect the human pathology and the use of human tumors faces several challenges such as ethical issues and the difficulty to access sufficient amount of fresh tissues to culture tumoroids, thus limiting high-throughput screening. In the present study, we have established the culture of canine mammary tumoroids in order to develop a biobank which could be used to provide a better comprehension of breast cancer pathogenesis and for large scale drug screening and therapeutic development for both veterinary and human medicine. In fact, dogs develop naturally numerous tumors in the presence of a functioning immune system that have similar features to human cancers(6,33–35). CMT are the most commonly diagnosed cancer in female dogs (50% of all cancers), which is a significant advantage since a large cohort of dogs could be recruited for preclinical studies.

Subtype classification of CMT has been investigated in a number of studies using IHC expression of PR, ER and HER2. Several distinct subtypes were identified including luminal A (14.3%), luminal B (9.4%) and triple negative (76.3%), while no HER2-overexpressing CMT were observed (7). Of the 6 dog patients included in the study, 4 tumors were of triple-negative subtype while 2 tumors were of luminal subtype, representing 67% of triple negative tumors and 33% of luminal tumors, which is consistent with previous findings. In human, as in dogs, the triple negative subtype is more aggressive leading to shorter survival rates compared to other tumors (33). Since therapeutic options for this subtype are limited, developing a reliable model to discover new effective treatments is highly needed.

We successfully generated tumoroids from CMT with a success rate of more than 94%. For comparison, in a recent study, human tumoroid establishment efficiency was around 40% for triple negative subtype(36). This difference can be explained by a higher amount of tissue which can be obtained from canine tumors, leading to high success rates. These tumoroids keep similar histological features as the original tumors as well as the same molecular subtype. Moreover, by a global proteomic analysis, we have shown that tumoroids were highly similar to the original tumor, 74% of proteins were identified in common between tumoroids and tumor. The tumor specific signature is of course due to a higher cellular diversity in the primary tumor compared to tumoroids, as demonstrated by the over-expression of immune related proteins such as proteins of the complement pathway. On the contrary, an enriched metabolic signature is noticed in tumoroids compared to the primary tumor, which can be explained by the stress of the culture triggering a higher cellular activity. Nevertheless, a high degree of similarity is kept between tumor and tumor-derived tumoroids. Interestingly, a recent study found that the main pathways that were enriched in breast cancer were linked to cell communication, cell growth and maintenance and signal transduction, which correlate well with our findings and is an additional proof that CMT are highly similar to human breast cancer(37).

Recently many tumoroid biobanks have emerged from different cancer types(38). Many of these studies have demonstrated that tumoroids preserve the genetic composition of the original tumor. However, the extent of molecular drift at later passage and after cryopreservation has been relatively low studied so far. In the presented study, we have compared histologic and molecular features (marker-based subtype and global proteome) as well as therapeutic response of tumoroids maintained in culture without cryopreservation (fresh tumoroids), put in culture after cryopreservation (frozen tumoroids) or developed from frozen cells (issued from the initial tumor, frozen cell tumoroids). We found that from a morphological point of view, the three types of tumoroids were similar and kept the same architecture and growth rates. The CMT subtype was also maintained after cryopreservation. We have also found that the type of culture or the number of passages did not impact too much the proteome of tumoroids. Indeed, the main variations were observed between tumoroids derived from different tumors rather than between different culture conditions. However, with this global proteomic analysis, we still found that fresh tumoroids and tumoroids made from frozen cells were more similar with a higher proteome diversity compared to frozen tumoroids. A previous study showed that tumor heterogeneity and cell diversity was conserved between fresh tumor tissue and cryopreserved tissue fragments or from cryopreserved cell suspensions (39). In the same study, the authors found that cryopreserved cell suspensions displayed higher correlations to fresh cells compared to tissue fragments. This can therefore explain our observations. Moreover, maintenance of stromal cell populations in tumoroids system is really challenging. At this time, the tumoroids culture system promote the expansion of the tumor cells but do not support the maintenance of immune cells and stromal cells(40). Stromal cells and immune cells are maintained during the first passages and tend to decrease overtime. By using an air-liquid interface to reconstitute the tumor microenvironment, tumoroids integrating immune components were successfully generated but immune cells tend to decline over time(41). Our proteomics results tend to demonstrate this fact, when tumoroids are kept fresh or are made from cells frozen after tumor dissociation, many proteins involved in metabolism, cell communication and immune response were identified. This immune signature was even much more pronounced for fresh tumoroids as demonstrated by the expression of T cell and macrophage markers (Granzyme B, Siglec1 and CD163). These results suggest that the cellular diversity may be higher in fresh tumoroids and tumoroids made from frozen cells compared to cryopreserved tumoroids. Metabolic and stress signatures were enriched in frozen tumoroids, which can be explained by cryopreservation(39).

To finish demonstrating that the culture conditions do not impact too much the tumoroids behavior, we have performed a drug response of tumoroids with a known chemotherapy used in human medicine. Paclitaxel response was similar between tumoroids, whatever the condition (fresh or cryopreserved). We however observed that cryopreserved tumoroids were slightly more resistant to Paclitaxel, reflected by a higher concentration of drug needed to kill 50% of cancer cells. These results corroborate our previous observations. Nevertheless, CMT tumoroids are sensitive to a human chemotherapy in a dose dependent manner with a similar response as human breast tumoroids.

## 5 Conclusions

In conclusion, for the first time, dog mammary tumoroids were produced from heterogeneous tumors. The tumoroids recapitulated the tumor histologic and molecular heterogeneities. Cryopreservation, which is often used for bio banking, did not seem to affect the molecular features and drug response of tumoroids. Nevertheless, we showed that cryopreservation of tumor cells after dissociation seem to best mimic the fresh tumoroids, with a higher molecular diversity. Canine tumoroids can be used to screen human drugs without limitations about tissue availability allowing large scale production. However, to make tumoroids even closer to the primary tumor, it is necessary to develop tumoroid models including stromal components such as immune cells which are lost during traditional tumoroids culture.

## Acknowledgements

The authors would like to thank Dr Alexandra Deviers, veterinary pathologist, for histopathological classifications of the canine mammary tumors used in the study and Dr Emmanuelle Cottin, veterinarian who provided us part of the tumor samples.

## Availability of data and materials

The mass spectrometry proteomics datasets generated and analyzed during the current study have been deposited to the ProteomeXchange Consortium via the PRIDE partner repository and are available with dataset identifier PXD031440.

## Authors’ contributions

ARR, MD and MS wrote the manuscript. MD and MS were responsible for the concept and design of the overall study and interpretation of the data. ARR and MD were involved in the experimental design, acquisition of the data and analysis and interpretation of the data. SA and EB provided technical assistance and guidance. NH, AR and DT provided the samples. MD, MS and IF supervised the project and provided critical revision of manuscript. MD, MS, IF and DT have obtained funding. All authors read and approved the final manuscript.

## Ethics approval and consent to participate

All animal and human studies were reviewed and approved by the local ethics committees, as detailed in the Materials and Methods section.

---

**<u>Supplementary Figure 1:</u>** Histology and receptor status (ER, PR, HER2) of the 6 canine mammary tumors used in the study.

**<u>Supplementary Figure 2:</u>** Immunofluorescence images of Ki67 stained canine mammary tumors.

**<u>Supplementary Table 1:</u>** Summary table of canine tumors used in the study.

**<u>Supplementary Table 2:</u>** List of proteins in the Venn diagram representing proteins identified in Fresh, Frozen, and FrozenCell tumoroids at date 1 and date 2.

**<u>Supplementary Table 3:</u>** List of proteins in the Venn diagram representing proteins in the Fresh, Frozen and FrozenCell tumoroids.

**<u>Supplementary Table 4:</u>** List of proteins in the 6 clusters identified in the Hierarchical clustering of the most variable proteins between the Fresh, Frozen and FrozenCell conditions.

**<u>Supplementary Table 5:</u>** List of proteins in the Venn diagram representing proteins in the Tumor, Fresh, Frozen and FrozenCell tumoroids conditions.

**<u>Supplementary Table 6:</u>** List of proteins in the 2 clusters identified in the Hierarchical clustering of the most variable proteins between the Tumor, Fresh, Frozen and FrozenCell tumoroids conditions.

## References

1. Wong CH, Siah KW, Lo AW. Estimation of clinical trial success rates and related parameters. Biostatistics [Internet]. 2019 [cited 2019 Oct 17];20(2):273–86. Available from: http://www.ncbi.nlm.nih.gov/pubmed/29394327

2. Broutier L, Mastrogiovanni G, Verstegen MM, Francies HE, Gavarró LM, Bradshaw CR, et al. Human primary liver cancer-derived organoid cultures for disease modeling and drug screening. Nat Med [Internet]. 2017 Dec [cited 2019 Sep 10];23(12):1424–35. Available from: http://www.ncbi.nlm.nih.gov/pubmed/29131160

3. Sachs N, de Ligt J, Kopper O, Gogola E, Bounova G, Weeber F, et al. A Living Biobank of Breast Cancer Organoids Captures Disease Heterogeneity. Cell. 2018 Jan 11;172(1–2):373–386.e10.

4. Nuciforo S, Fofana I, Matter MS, Blumer T, Calabrese D, Boldanova T, et al. Organoid Models of Human Liver Cancers Derived from Tumor Needle Biopsies. Cell Rep [Internet]. 2018 Jul 31 [cited 2019 Sep 10];24(5):1363–76. Available from: http://www.ncbi.nlm.nih.gov/pubmed/30067989

5. Lee SH, Hu W, Matulay JT, Silva M V., Owczarek TB, Kim K, et al. Tumor Evolution and Drug Response in Patient-Derived Organoid Models of Bladder Cancer. Cell. 2018 Apr 5;173(2):515–528.e17.

6. Nguyen F, Peña L, Ibisch C, Loussouarn D, Gama A, Rieder N, et al. Canine invasive mammary carcinomas as models of human breast cancer. Part 1: natural history and prognostic factors. Breast Cancer Res Treat [Internet]. 2018 [cited 2019 Oct 17];167(3):635–48. Available from: http://www.ncbi.nlm.nih.gov/pubmed/29086231

7. Abadie J, Nguyen F, Loussouarn D, Peña L, Gama A, Rieder N, et al. Canine invasive mammary carcinomas as models of human breast cancer. Part 2: immunophenotypes and prognostic significance. Breast Cancer Res Treat [Internet]. 2018 [cited 2019 Oct 17];167(2):459–68. Available from: http://www.ncbi.nlm.nih.gov/pubmed/29063312

8. Dorn CR, Taylor DON, Schneider R, Hibbard HH, Klauber MR. Survey of animal neoplasms in alameda and contra costa counties, california. ii. cancer morbidity in dogs and cats from alameda county. J Natl Cancer Inst. 1968;40(2):307–18.

9. Dobson JM, Samuel S, Milstein H, Rogers K, Wood JLN. Canine neoplasia in the UK: estimates of incidence rates from a population of insured dogs. J Small Anim Pract [Internet]. 2002 Jun [cited 2019 Oct 17];43(6):240–6. Available from: http://www.ncbi.nlm.nih.gov/pubmed/12074288

10. Egenvall A, Bonnett BN, Ohagen P, Olson P, Hedhammar A, von Euler H. Incidence of and survival after mammary tumors in a population of over 80,000 insured female dogs in Sweden from 1995 to 2002. Prev Vet Med [Internet]. 2005 Jun 10 [cited 2019 Oct 17];69(1–2):109–27. Available from: http://www.ncbi.nlm.nih.gov/pubmed/15899300

11. MacEwen EG. Spontaneous tumors in dogs and cats: Models for the study of cancer biology and treatment. CANCER METASTASIS Rev. 1990 Sep;9(2):125–36.

12. Queiroga FL, Raposo T, Carvalho MI, Prada J, Pires I. Canine mammary tumours as a model to study human breast cancer: Most recent findings. Vol. 25, In Vivo. 2011. p. 455–65.

13. Michałowska M, Winiarczyk S, Adaszek Ł, Łopuszyński W, Grądzki Z, Salmons B, et al. Phase I/II clinical trial of encapsulated, cytochrome P450 expressing cells as local activators of cyclophosphamide to treat spontaneous canine tumours. PLoS One [Internet]. 2014 [cited 2019 Oct 17];9(7):e102061. Available from: http://www.ncbi.nlm.nih.gov/pubmed/25028963

14. T U, M S, S N, K U, Y N, T H, et al. Establishment of a dog primary prostate cancer organoid using the urine cancer stem cells. Cancer Sci [Internet]. 2017 Dec 1 [cited 2021 Nov 2];108(12):2383–92. Available from: https://pubmed.ncbi.nlm.nih.gov/29024204/

15. Kramer N, Pratscher B, Meneses AMC, Tschulenk W, Walter I, Swoboda A, et al. Generation of Differentiating and Long-Living Intestinal Organoids Reflecting the Cellular Diversity of Canine Intestine. Cells 2020, Vol 9, Page 822 [Internet]. 2020 Mar 28 [cited 2021 Nov 2];9(4):822. Available from: https://www.mdpi.com/2073-4409/9/4/822/htm

16. Ambrosini YM, Park Y, Jergens AE, Shin W, Min S, Atherly T, et al. Recapitulation of the accessible interface of biopsy-derived canine intestinal organoids to study epithelial-luminal interactions. PLoS One [Internet]. 2020 Apr 1 [cited 2021 Nov 2];15(4):e0231423. Available from: https://journals.plos.org/plosone/article?id=10.1371/journal.pone.0231423

17. Chandra L, Borcherding DC, Kingsbury D, Atherly T, Ambrosini YM, Bourgois-Mochel A, et al. Derivation of adult canine intestinal organoids for translational research in gastroenterology. BMC Biol [Internet]. 2019 Apr 11 [cited 2021 Nov 2];17(1). Available from: /pmc/articles/PMC6460554/

18. C C, S M, E P, MC V, M G, C B, et al. FGF2 and EGF Are Required for Self-Renewal and Organoid Formation of Canine Normal and Tumor Breast Stem Cells. J Cell Biochem [Internet]. 2017 Mar 1 [cited 2021 Nov 2];118(3):570–84. Available from: https://pubmed.ncbi.nlm.nih.gov/27632571/

19. Botti G, Di Bonito M, Cantile M. Organoid biobanks as a new tool for pre-clinical validation of candidate drug efficacy and safety. Int J Physiol Pathophysiol Pharmacol [Internet]. 2021 [cited 2021 Nov 2];13(1):17–21. Available from: https://clinicaltrials.

20. Wisniewski JR, Zougman A, Nagaraj N, Mann M. Universal sample preparation method for proteome analysis. Nat Methods. 2009 May;6(5):359–62.

21. Cox J, Mann M. MaxQuant enables high peptide identification rates, individualized p.p.b.-range mass accuracies and proteome-wide protein quantification. Nat Biotechnol 2008 2612. 2008 Nov;26(12):1367–72.

22. Cox J, Neuhauser N, Michalski A, Scheltema RA, Olsen J V., Mann M. Andromeda: A peptide search engine integrated into the MaxQuant environment. J Proteome Res. 2011;10(4):1794–805.

23. Tyanova S, Cox J. Perseus: A Bioinformatics Platform for Integrative Analysis of Proteomics Data in Cancer Research. Methods Mol Biol. 2018;1711:133–48.

24. Tyanova S, Temu T, Sinitcyn P, Carlson A, Hein MY, Geiger T, et al. The Perseus computational platform for comprehensive analysis of (prote)omics data. Nat Methods 2016 139. 2016 Jun;13(9):731–40.

25. Mi H, Huang X, Muruganujan A, Tang H, Mills C, Kang D, et al. PANTHER version 11: expanded annotation data from Gene Ontology and Reactome pathways, and data analysis tool enhancements. Nucleic Acids Res. 2017 Jan;45(D1):D183–9.

26. Szklarczyk D, Franceschini A, Wyder S, Forslund K, Heller D, Huerta-Cepas J, et al. STRING v10: Protein-protein interaction networks, integrated over the tree of life. Nucleic Acids Res. 2015 Jan;43(D1):D447–52.

27. Pathan M, Keerthikumar S, Ang CS, Gangoda L, Quek CYJ, Williamson NA, et al. FunRich: An open access standalone functional enrichment and interaction network analysis tool. Proteomics. 2015 Aug;15(15):2597–601.

28. Goldschmidt M, Peña L, Rasotto R, Zappulli V. Classification and grading of canine mammary tumors. Vet Pathol [Internet]. 2011 Jan [cited 2019 Oct 22];48(1):117–31. Available from: http://www.ncbi.nlm.nih.gov/pubmed/21266722

29. Song P, Yang F, Jin H, Wang X. The regulation of protein translation and its implications for cancer. Signal Transduct Target Ther 2021 61 [Internet]. 2021 Feb 18 [cited 2022 Jan 31];6(1):1–9. Available from: https://www.nature.com/articles/s41392-020-00444-9

30. Imamura T, Yamamoto-Ibusuki M, Sueta A, Kubo T, Irie A, Kikuchi K, et al. Influence of the C5a–C5a receptor system on breast cancer progression and patient prognosis. Breast Cancer 2015 236 [Internet]. 2015 Oct 22 [cited 2022 Jan 31];23(6):876–85. Available from: https://link.springer.com/article/10.1007/s12282-015-0654-3

31. Boyer DS, Rippmann JF, Ehrlich MS, Bakker RA, Chong V, Nguyen QD. Amine oxidase copper-containing 3 (AOC3) inhibition: a potential novel target for the management of diabetic retinopathy. Int J Retin Vitr [Internet]. 2021 Dec 1 [cited 2022 Jan 31];7(1). Available from: https://pubmed.ncbi.nlm.nih.gov/33845913/

32. Mohan SC, Lee TY, Giuliano AE, Cui X. Current Status of Breast Organoid Models. Front Bioeng Biotechnol. 2021 Nov 5;9:1091.

33. Gray M, Meehan J, Martínez-Pérez C, Kay C, Turnbull AK, Morrison LR, et al. Naturally-Occurring Canine Mammary Tumors as a Translational Model for Human Breast Cancer. Front Oncol. 2020 Apr 28;10:617.

34. Uva P, Aurisicchio L, Watters J, Loboda A, Kulkarni A, Castle J, et al. Comparative expression pathway analysis of human and canine mammary tumors. BMC Genomics. 2009 Mar 27;10.

35. Abdelmegeed SM, Mohammed S. Canine mammary tumors as a model for human disease (Review). Vol. 15, Oncology Letters. Spandidos Publications; 2018. p. 8195–205.

36. Dekkers JF, van Vliet EJ, Sachs N, Rosenbluth JM, Kopper O, Rebel HG, et al. Long-term culture, genetic manipulation and xenotransplantation of human normal and breast cancer organoids. Nat Protoc 2021 164 [Internet]. 2021 Mar 10 [cited 2022 Jan 25];16(4):1936–65. Available from: https://www.nature.com/articles/s41596-020-00474-1

37. Deng JL, Xu YH, Wang G. Identification of potential crucial genes and key pathways in breast cancer using bioinformatic analysis. Front Genet. 2019;10(JUL):695.

38. Lo YH, Karlsson K, Kuo CJ. Applications of Organoids for Cancer Biology and Precision Medicine. Nat cancer [Internet]. 2020 Aug 1 [cited 2022 Jan 25];1(8):761. Available from: /pmc/articles/PMC8208643/

39. Wu SZ, Roden DL, Al-Eryani G, Bartonicek N, Harvey K, Cazet AS, et al. Cryopreservation of human cancers conserves tumour heterogeneity for single-cell multi-omics analysis. Genome Med [Internet]. 2021 Dec 1 [cited 2022 Jan 26];13(1):1–17. Available from: https://genomemedicine.biomedcentral.com/articles/10.1186/s13073-021-00885-z

40. Fiorini E, Veghini L, Corbo V. Modeling Cell Communication in Cancer With Organoids: Making the Complex Simple. Front Cell Dev Biol. 2020 Mar 18;8:166.

41. Neal JT, Li X, Zhu J, Giangarra V, Grzeskowiak CL, Ju J, et al. Organoid Modeling of the Tumor Immune Microenvironment. Cell. 2018 Dec 13;175(7):1972–1988.e16.

