## Supplementary figures and images for "Establishment and characterization of a tumoroid biobank derived from dog patients’ mammary tumors for translational research"

### Supplementary Figure 1

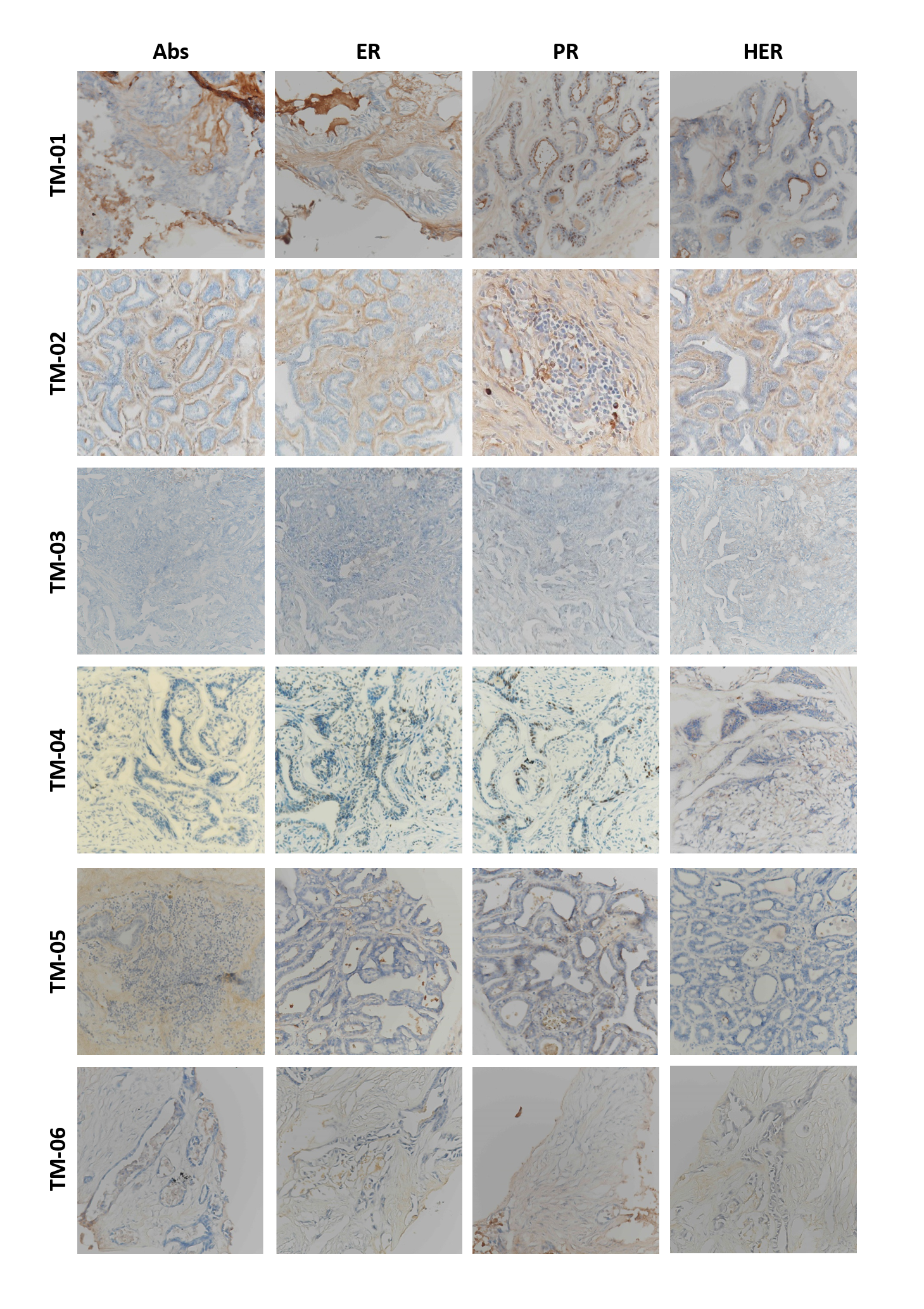

### Supplementary Figure 2

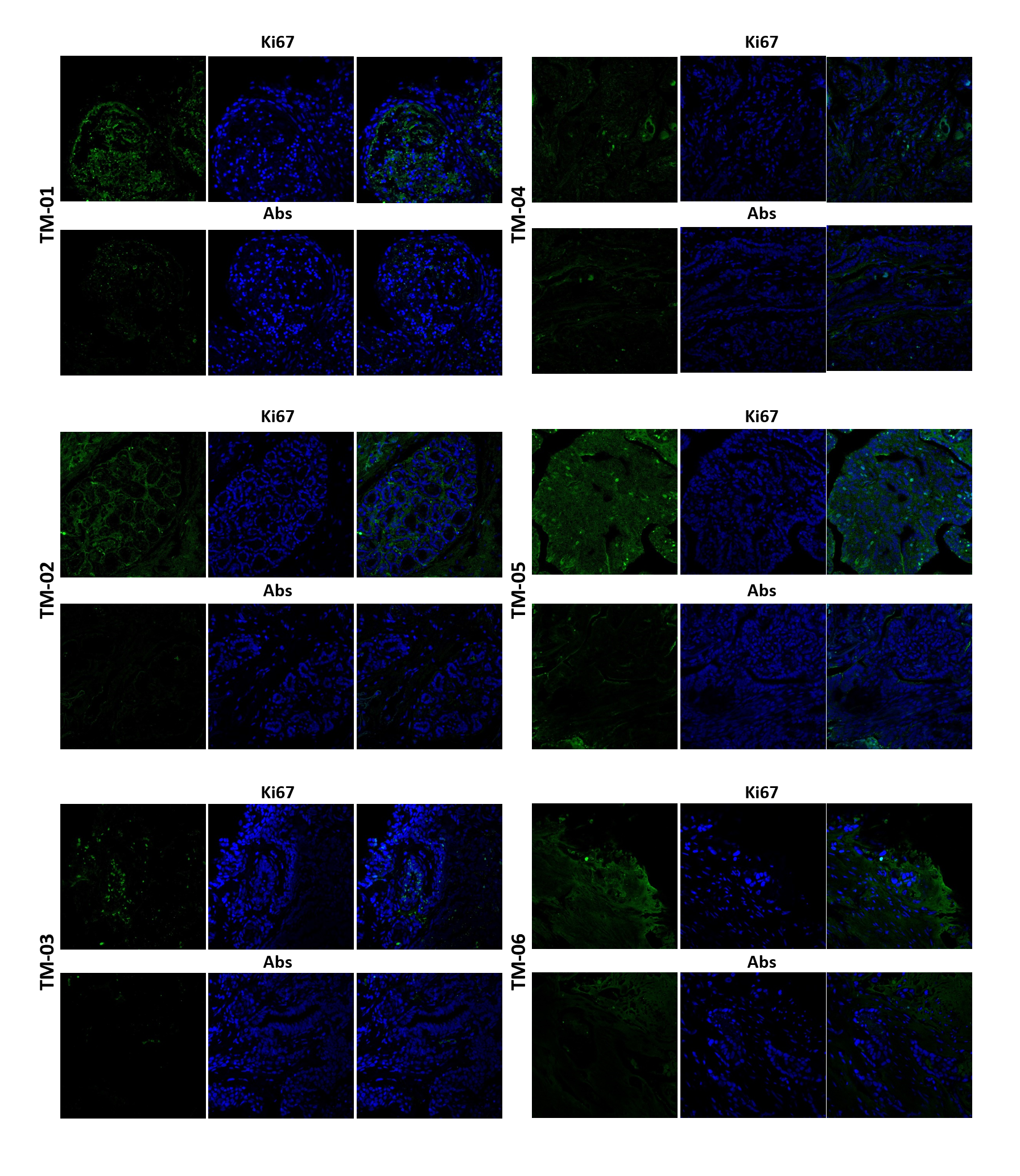
